# Pollen metabarcoding as a tool for tracking long-distance insect migrations

**DOI:** 10.1101/312363

**Authors:** Tomasz Suchan, Gerard Talavera, Llorenç Sáez, Michał Ronikier, Roger Vila

## Abstract

Insects account for the main fraction of Earth’s biodiversity and are key players for ecosystems, notably as pollinators. While insect migration is suspected to represent a natural phenomenon of major importance, remarkably little is known about it, except for a few flagship species. The reason for this situation is mainly due to technical limitations in the study of insect movement. Here we propose using metabarcoding of pollen carried by insects as a method for tracking their migrations. We developed a flexible and simple protocol allowing high multiplexing and not requiring DNA extraction, one of the most time consuming part of metabarcoding protocols, and apply this method to the study of the longdistance migration of the butterfly *Vanessa cardui,* an emerging model for insect migration. We collected 47 butterfly samples along the Mediterranean coast of Spain in spring and performed metabarcoding of pollen collected from their bodies to test for potential arrivals from the African continent. In total, we detected 157 plant species from 23 orders, most of which (82.8%) were insect-pollinated. African or African-Arabian endemic taxa contributed 21.0% of our dataset, strongly supporting the hypothesis that migratory butterflies colonize southern Europe from Africa in spring. Moreover, our data suggest that a northwards trans-Saharan migration in spring is plausible for early arrivals (February) into Europe, as shown by the presence of Saharan floristic elements. Our results demonstrate the possibility of regular insect-mediated transcontinental pollination, with potential implications for ecosystem functioning, agriculture and plant phylogeography. Despite current limitations, mostly regarding the availability of plant reference sequences and distribution data, the method proved to be useful and demonstrates great potential as plant genetic libraries and distribution datasets improve.

## Introduction

Insects undergo aerial long-distance migrations (Holland, Wikielski & Wilcove, 2006; Chapman, Reynolds & Wilson, 2015) that outnumber migrations of larger organisms, such as birds, both in abundance and biomass (Hu et al., 2016). These long-range movements have important –albeit still largely unknown– implications for ecosystems and human economy (Bauer & Hoye, 2014; Chapman et al., 2015; Hu et al., 2016). Nevertheless, mostly due to the technical challenges associated with tracking small organisms (Chapman et al., 2015), our knowledge on insect migrations is extremely limited (Holland et al., 2006), especially in comparison to that on vertebrate migrations.

Tracking long-distance insect migrations involves assessing the actual path of individuals, either by mark-recapture studies, using variety of markers (reviewed in Hagler & Jackson, 2001), or by telemetry (Osborne, Loxdale & Woiwod, 2002; Kissling, Pattemore & Hagen, 2014). However, mark-recapture studies of migrating insects have low recapture success rate and are feasible only for the most emblematic species, such as monarch butterflies (Knight, Brower & Williams, 1999; Garland & Davis, 2002). On the other hand, radio telemetry is only suitable for tracking the largest insects at short distances, and is technically limited by relatively high weight of the transmitters and short battery life (Kissling et al., 2014). Because of that, so far only one telemetric study was performed to assess long-distance migrations in insects – of the dragonfly *Anax junius.* It however involved considerable logistic challenges, such as using small planes and ground teams for tracking insect movements (Wikelski et al., 2006).

Due to the above limitations, insect migration studies are traditionally observation-based, linking the place, time and, sometimes, direction of the observations with hypothetical routes (e.g. Howard & Davis, 2009; Talavera & Vila, 2016). More recently, other observational methods, such as vertical-looking radars (Chapman, Drake & Reynolds, 2011) or Doppler weather radars (Chilson et al., 2012; Westbrook & Eyster, 2017) have been used in insect migration studies, often supplemented by aerial nets or other type of traps for accurate species determination (Chapman et al., 2002; Chapman et al., 2004). The new developments in this technology allow insect body mass, shape, wing-beat frequency, flight altitude and heading directions to be measured, often allowing species determination (Dean & Drake, 2005; Chapman et al., 2011; Drake et al. 2017), but observations are usually constrained to particular areas because radars have no or limited mobility.

Other methods of studying migrations use intrinsic markers, such as genetic markers or isotope composition. By using genetic markers, populations connected by regular gene flow can be identified (Lowe & Allendorf, 2010), which may suggest migratory routes and natural barriers (e.g. Nagoshi, Meagher & Hay-Roe, 2012). The utility of genetic markers is dependent on spatial genetic structuring. For migratory insects, structuring is expected between independent migratory circuits, but might not be maintained in the case of regular gene flow between migrating lineages (global panmixia) (Lyons et al., 2012; Lukhtanov, Pazhenkova & Novikova 2016). Because the stable isotope ratios, such as ^2^H/^1^H or ^13^C/^12^C, of organic tissues are related to the site where insects developed, these can also be used, along with modelled geographic isotope patterns (isoscapes), to infer probabilistic natal origins of migrating individuals (e.g. Hobson, Wassenaar & Taylor, 1999; Brättstrom et al., 2010; Stefanescu, Soto, Talavera, Vila & Hobson, 2016). This technique does not rely on marking-recapturing specimens and it is thus also suitable for small species (Hobson, 2008). However geospatial assignments depend on limited isoscape resolution and are usually only helpful at inferring large-scale geographic patterns.

As insects visit and feed on flowers, the pollen is deposited on their bodies and can be transported across large distances (Ahmed et al., 2009). Therefore, pollen of plants endemic to certain areas could also be exploited for tracking long-distance insect migrations (Hagler & Jackson, 2001; Jones & Jones, 2001) and, indeed, it has been used as a marker in a handful of studies (Mikkola, 1971; Hendrix, Mueller, Phillips & Davis, 1987; Hendrix & Showers, 1992; Gregg, 1993; Lingren et al., 1993, 1994; Westbrook et al., 1997). However, conventional pollen identification by light or electron microscopy is time-consuming, and requires specialized taxonomic knowledge. It is therefore difficult to apply as a widely accessible tool for large-scale studies (Galimberti et al., 2014; Keller et al., 2015; Richardson et al., 2015b; Sickel et al., 2015). Moreover, taxonomical classification of pollen grains is often unachievable to the species or even genus level (Hawkins et al., 2015; Kraaijeveld et al., 2015; Richardson et al., 2015b).

The development of the next generation sequencing (NGS) technologies allowed straightforward sequencing of DNA barcodes from mixed environmental samples, termed “metabarcoding” (Taberlet et al., 2012; Deiner et al., 2017). In this study, we use a DNA metabarcoding approach to identify pollen grains carried by a long-distance migratory insect species – the painted lady butterfly *Vanessa cardui.* This is a virtually cosmopolitan species adapted to seasonally exploit a wide range of habitats and sometimes observed even at extreme latitudes or in the open ocean (Shields, 1992). The Palearctic-African migratory system involves populations that undergo yearly long-distance latitudinal migrations in a circuit between Tropical Africa (September to February) and the temperate zone (February to September) (Talavera & Vila, 2016). Such annual circuit might involve 10 generations or more, some of them performing long-distance movements of thousands of kilometres. The distances crossed by individual butterflies within a single generation are unclear. It has been recently shown, both by stable isotope and observational evidence, that the species massively immigrate and breed in autumn in sub-Saharan Africa, and that these populations are of European origin (Stefanescu, Soto, Talavera, Vila & Hobson, 2016; Talavera & Vila, 2016). These results depict a new spatiotemporal model for the migration of *V. cardui* in this part of the world, which involves migratory movements across differentiated floristic regions: central and northern Europe, the Mediterranean, the Sahara and Tropical Africa. Although the phenology of migratory movements within the western Palearctic is well known, both from observations and entomological radars (Stefanescu et al., 2013), the African locales and routes of the species from October to March are unclear (Talavera & Vila, 2016).

Using pollen metabarcoding from captured butterflies apparently migrating northwards into Europe, we test i) for the presence of DNA from pollen grains deposited on insects’ bodies after a long-distance migration from Africa and, if detected, ii) whether the sequences obtained are of African endemic plant species that could be informative on the migration routes.

Despite great potential of pollen DNA metabarcoding for pollination biology or palynological studies, it has been used only in a handful of cases, mostly to investigate honey composition (Valentini, Miquel & Taberlet, 2010; Bruni et al., 2015; Hawkins et al., 2015; Prosser & Herbert, 2017), honeybee foraging (Galimberti et al., 2014; Richardson et al., 2015a, b; de Vere, 2017) and, recently, plant-pollinator interactions (Keller et al., 2015; Sickel et al., 2015; Pornon et al., 2016; Bell et al., 2017). Several DNA markers have been proposed for identifying mixed pollen loads from insects (Bell et al., 2016), the Internal Transcribed Spacer 2 (ITS2) nuclear ribosomal fragment being one of the most frequently employed (Keller et al., 2015; Richardson et al., 2015b; Sickel et al., 2015). ITS2 has several advantages for pollen metabarcoding. Although plastid markers have been successfully amplified from pollen grains (e.g. Kraaijeveld et al., 2015; Richardson et al., 2015a), the variation in the number of plastid genome copies is poorly understood (Bell et al., 2016). Because it is a nuclear marker, ITS2 should be more uniformly present in pollen. Moreover, the sequence length of this marker is short enough for amplicon sequencing using Illumina MiSeq technology. It is also sufficiently informative to discriminate most plant species (Chen et al., 2010) and the number of available reference sequences in databases is the highest among the available plant barcodes (Bell et al., 2016).

Several library preparation protocols have been proposed for pollen metabarcoding. Richardson et al. (2015b) performed ITS2 amplification using standard primers followed by purification and NGS library preparation using commercial kits. Keller et al. (2015) and Sickel et al. (2015) proposed an approach based on Kozich, Westcott, Baxter, Highlander and Schloss (2013) protocol, where an ITS2 amplicon Illumina library is prepared within a single PCR step. These authors used ITS2 primers tailed with appropriate technical sequences and custom sequencing primers, to avoid losing sequencing cycles for ITS2 primer sequences and avoid problems related to low sequence diversity of these regions. The downside of such protocol is the need of replacing all the oligonucleotides when targeting other markers, template-specific biases when using indexed primers (O’Donnell, Kelly, Lowell & Port, 2016), and the necessity to use custom sequencing primers.

Here, we developed a laboratory protocol with maximum flexibility, cost-effectiveness and reduced workload in mind – especially by lowering the number of library preparation steps, which not only require more work, but can also lead to sample cross-contamination. Our method consists of two PCR steps: first, the ITS2 fragment is amplified using standard primers (White, Bruns, Lee & Taylor, 1990; Chen et al., 2010), tailed with partial Illumina sequences; in the second step, the fragments are double-indexed and the final library is produced – an approach similar to the Illumina 16S sequencing protocol (http://res.illumina.com/documents/products/appnotes/appnote_miseq_16s.pdf). Compared to the protocol by Sickel et al. (2015), our approach uses one additional PCR reaction, but it is more flexible and cost-effective because the sequenced DNA barcode can be changed just by modifying the two primers used in the first PCR reaction. Moreover, in the second PCR reaction we use primers compatible with standard Illumina sequencing protocol, which simplifies the sequencing step and allows sequencing the libraries along with other standard libraries. We also contribute a bioinformatic pipeline to analyse and classify the obtained reads.

## Materials and methods

### Sampling

We monitored seven sites along the Mediterranean coast of Spain to test for potential arrivals of *Vanessa cardui* from the African continent (Fig. 1, Tab. S1). Sampling was designed to capture specimens with high probability to be in migratory phase. To do that, we sampled sites where the species was unlikely to be found while nectaring or breeding. In particular we sampled points in the Mediterranean shorelines, usually in the sandy beaches, cliffs or inspecting the vegetation nearby the coast. Timing was also chosen within the time frame where *V. cardui* arrivals are expected to colonize the Iberian Peninsula (February-April), and when consistent wind patterns or storms from Africa occurred, that could aid insect migrations. All the samples were immediately bagged in glassine envelopes that were sealed and stored at −20°C until pollen isolation and library preparation.

**Fig. 1.**
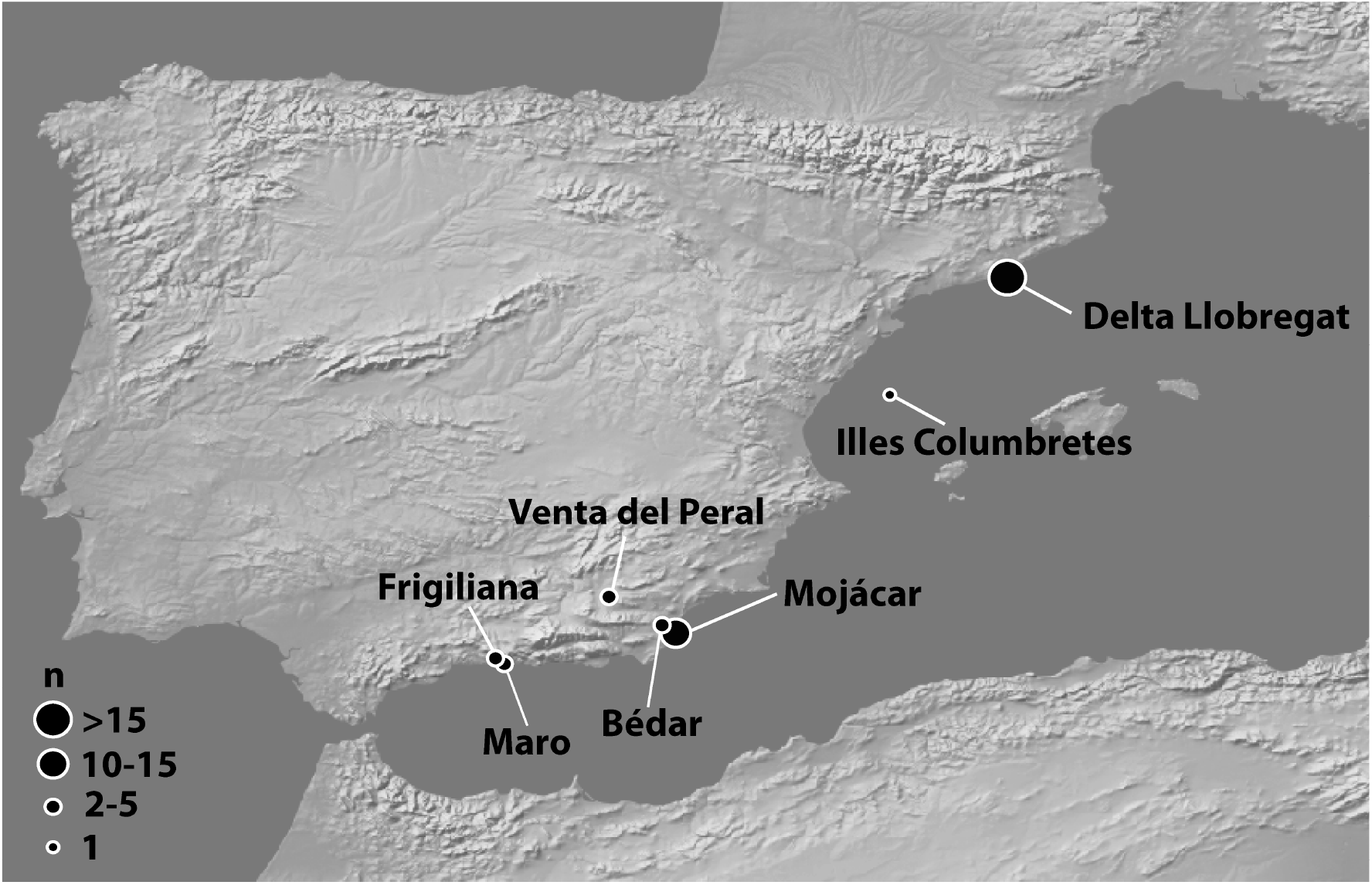
Map of the sampled specimens. The size of the points indicates the number of samples collected in each locality.

### NGS library construction

Pollen isolation and library construction were performed in four batches (see Tab. S2). We amplified the Internal Transcribed Region 2 (ITS2) of the ribosomal DNA by using a combination of ITS-S2F (Chen et al., 2010) and ITS-4R (White et al., 1990) primers, tested in other pollen metabarcoding studies (Keller et al., 2015; Sickel et al., 2015). In the first step, the ITS2 fragment was amplified using the above primers tailed with technical sequences: six random nucleotides to increase sequence diversity during the first sequencing cycles, and a part of the Illumina adapter. In the second reaction, we used modified Illumina TruSeq primers (Tab. 1) in order to index the samples and produce the final library. We used index sequences from TruSeq Amplicon series, allowing the pool of 96 samples on one lane. These indexes can be supplemented by TruSeq CD index sequences (see https://support.illumina.com/content/dam/illumina-support/documents/documentation/chemistry_documentation/experiment-design/illumina-adapter-sequences-1000000002694-03.pdf) as in Sickel et al. (2015), for a total of 384 combinations. All the steps of the protocol, from pollen isolation up to the first PCR reaction were conducted under a laminar flow cabinet in a pre-PCR area in order to avoid external contamination. All the working spaces and equipment were cleaned with 10% bleach solution and 70% ethanol before and after work. We used only ultra-pure DEPC-treated water and a separate stock of reagents and plastics dedicated solely to metabarcoding work. Moreover, library preparation steps for all the samples and pollen isolation from most of the samples (batches 1-3) were conducted in a laboratory located far away from the sampling sites (at W. Szafer Institute of Botany, Polish Academy of Sciences, Kraków, Poland). Pollen isolation from batch 4 was performed at Institut de Biologia Evolutiva, Barcelona, Spain.

After preliminary trials (not shown), we decided not to extract DNA but to use Phire Plant Direct Polymerase (Thermo Fisher Scientific, Waltham, MT, USA) in the first PCR step, which successfully amplified ITS2 from pollen mix without DNA extraction. Pollen was collected by vortexing butterfly bodies (with wings removed) in a 2 ml tube with 50 μl of sterile water with 0.1% SDS, centrifugation of the obtained solution and drying it under vacuum. The obtained pellet was diluted in 15 μl of Phire Plant Direct sample buffer and homogenized with five zirconium beads on the TissueLyser II machine (Qiagen, Hilden, Germany) at 30 Hz for 1 min before proceeding to the PCR. We added blank sample to each of the four extraction batches (“extraction blank samples”) – a 1.5 ml tube filled with 50 μl of water, left open during the whole extraction procedure and processed like a normal sample. We processed each sample in three independent PCR reactions to avoid reaction-specific biases (Fierer, Hamady, Lauber & Knight, 2008; Sickel et al., 2015) with another four blank samples added at the PCR step (“PCR blank samples”). Each of the replicate PCR reactions consisted of 1 μl of the disrupted pollen sample, 25 μl of Phire Plant Direct Polymerase Mix, and 0.5 μM of each primer in 50 μl reaction volume with the following PCR program: 98°C for 5 min, 20 cycles of denaturation at 98°C for 40 s, annealing at 49°C for 40 s and elongation at 72°C for 40 s, final extension step at 72°C for 5 min. After the reaction, we combined the PCR triplicates and purified the product using 1x ratio of AMPure XP (Beckman Coulter, Indianapolis, IN, USA) and eluted in 10 μl of water. In the second reaction, we indexed each sample using a unique combination of primers. The second reaction, also performed in triplicates for each sample, consisted of 1 μl of the purified PCR product, 1x Q5 buffer, 0.2 U of Q5 Hot-Start polymerase, 0.5 μM of forward, and 0.5 μM of reverse indexed primer in 10 μl reaction volume. We amplified the reaction using a PCR program as follows: 30 s initial denaturation at 98°C; 12 cycles of denaturation at 98°C for 10 s and combined annealing and extension at 72°C for 30 s (shuttle PCR); final extension at 72°C for 5 min. After the reaction, we combined the triplicates and verified the reaction success on TapeStation 4200 (Agilent, Santa Clara, CA, USA). We then pooled 10 μl of the PCR product from each sample and purified the pool using 1x ratio of AMPure XP. After that, the pooled library was quantified using the Qubit instrument (Thermo Fisher Scientific, Waltham, Massachusetts, USA), and sequenced with 15% PhiX spike-in on Illumina MiSeq (San Diego, CA, USA) using 600-cycle MiSeq Reagent Kit v3, according to the manufacturer’s instructions.

### Data analysis

We merged the raw paired-end reads using PEAR v0.9.8 (Zhang, Kobert, Flouri & Stamatakis, 2014) and retained only the successfully merged reads. We then trimmed the primer sequences with remove_primers script (www.biopieces.org). The obtained sequences were processed using vsearch v2.4.3 (Rognes, Flouri, Nichols, Quince & Mahé, 2016): filtered by an expected error rate *(maxe* parameter) of 0.5, minimum length of 250 nt, maximum length of 450 nt, and removing the reads with ambiguous nucleotides. Next, the reads were dereplicated and singletons removed. We classified the filtered reads with SINTAX (from the USEARCH package v10.0.240; Edgar, 2016a) using the ITS2 database of Sickel et al. (2016), modified for SINTAX by parsing with a custom script. Only reads classified to the species level with a probability of 95% or above were further considered. The classified reads were then summarized by species with a minimum threshold of 100 reads per plant species per sample. The raw sequences, the scripts used to process the data, and ITS2 database are available online (see Data Availability).

### Plant distributions

Geographical distributions of all detected plants were compiled checking specific literature and occurrences available in the online databases GBIF (www.gbif.org) and the African Plant Database (www.ville-ge.ch/musinfo/bd/cjb/africa/recherche.php) (Supplementary Table S1). The plants with the geographical range not including the butterfly sampling sites were further examined, as potential marker candidates to unravel migratory paths. These mainly included Saharan-North African endemics and African species with ranges reaching southern Mediterranean (Figure S1). The extent of occurrence for each informative plant species was delimited on the maps by the minimum convex polygon containing all known sites of occurrence, according to IUCN criteria (IUCN, 2012).

### Wind track

Backward wind trajectories were reconstructed for sampling sites and dates, using the Hybrid Single-Particle Lagrangian Integrated Trajectory (HYSPLIT) dispersion model from NOAA’s Air Resources Laboratory (ARL) (Stein et al., 2015; Rolph, Stein & Stunder, 2017). Analyses were based on the Reanalysis database and computed on 48-h back trajectories arriving at sites at 12:00 h UTC and for three different altitudinal layers (500m, 1000m and 1500m asl).

## Results

### Sampling

A total of 47 *V. cardui* specimens were collected in the seven sites (Fig. 1; Tab. S1) and processed for pollen metabarcoding analysis. The sampling consisted mostly of two migratory groups, according to the regions (Andalusia and Catalonia) and timing (February and April). First, a group of samples was collected in several localities of Andalusia between February 16^th^ and 25^th^ of 2016, a year with exceptionally early sightings of the species in the region.

The individuals from this group were generally collected close to the beach, but not exactly at the moment of arrival, although a noticeable increase of individuals was observed on the 21^st^ of February. A second group of samples were collected at the beach of Delta del Llobregat, near Barcelona, along two consecutive days (27^th^ and 28^th^) in April of 2012, after a storm carrying Saharan dust and strong winds from the south. The individuals from this group were collected when landing on the beach, coming from the sea in a northwest direction. Finally, one sample was collected in June 2012 from Columbretes Islands, a small archipelago 50 km from Castelló coast.

### Sequencing results and reads processing

We obtained 9,654,286 raw paired-end reads, of which only 1420 (0.0147%) were present in the blank samples. The average number of reads per sample was 205,380 (excluding blank samples; range: 7-608,640), of which 98.7% were successfully merged (Tab. S2). After quality filtering, we retained a total of 7,315,458 reads remained, of which 1,236,996 (16.9%) were discarded as singletons. The rest of the reads were dereplicated (separately within each sample) into 229,295 unique sequences of which 35,393 (15.4%; that is 2,908,006 reads or 47.8% of all the filtered reads) were classified to the species level with probability ≥ 95%. The proportion of the filtered reads classified to higher taxonomic ranks was much higher: 99.9% to the division, 78.3% to the order, and 65.7% to the family level (Fig. S2). After filtering out the plant species represented by less than 100 reads in a butterfly sample, 2,880,443 reads were retained (47.4% of the filtered reads; Tab. S2).

Sequencing success was not even among the butterfly samples but rather followed a bimodal distribution (Fig. S1a) – 17 samples yielded less than 10,000 raw sequences (mean = 1,041; range: 7-7,702) but the remaining 30 samples had high number of reads (mean = 321,172; range: 37,420-608,640; Tab S2). We also detected variation among the four sample batches: samples from batches number 1 and 3 had a low coverage, as did five out of 11 samples from batch 2. In contrast, most of the samples of batch 4 had a high coverage (Fig. S1b; Tab. S2). Sequences assigned to the species level with high probability and coverage higher than 100 reads were found in 30 out of 47 (63.8%) butterfly samples, from all the batches except number one but only in the samples with high the number of reads higher than 10,000. No reads were retained after the filtering steps for the sample from Columbretes Islands (the only one not attributable to Andalusia or Catalonia).

The amount of PCR product in blank samples was too low to be visible in TapeStation profiles. During the sequencing, PCR blank samples yielded from 0 to 5 reads. Extraction blank samples had between 3 and 26 raw reads, except EB4 from which we sequenced 1,366 reads. After processing the latter, we detected 42 unique sequences, of which only three were classified to the species level with probability ≥ 95%, belonging to *Moricandia moricandioides* (11 reads in total) and *Prunus dulcis* (27 reads). These two plant species were detected in some of the studied butterflies and are not present in the area where the libraries were prepared (Kraków, Poland), pointing rather to a low level of cross-contamination than to external contamination in one of the batches. As these sequences were well below the threshold of 100 reads per species, they were not retained in the final dataset.

### Sequence classification and plant diversity

In total, the filtered reads were classified to 157 species (Table S3). The sequence most frequently represented in the samples (present in 21 samples) was classified as *Alternanthera* sp. This appears to be an erroneous GenBank entry (JX136744.1), most likely of fungal origin and it was removed from our dataset. Excluding this, the most represented species (in 12 out of 30 butterfly samples for which plant sequences were detected) was *Reseda lutea* a widespread species in Europe and the Mediterranean region, including both sampling areas (Andalusia and Catalonia). Approximately half of the plant species (76) were present only in a single butterfly sample each and another 42 species in just two butterfly samples each. On average, there were 12.2 species detected per butterfly sample (SD: 6.9; range: 2-26).

The diversity of detected pollen included species from 23 orders of plants in total, where 13 orders were represented in samples from Catalonia (April) and 23 orders in Andalusia (February). Asterales were the dominant order at both sites, with 31 species: 14 and 21 species in the two above regions, respectively. Most of the plants detected were typically insect-pollinated (82.8%), while 15.3% were species not pollinated by insects. The proportion of insect pollinated species was higher at both sites: 78.7% Catalonia and 85.6% in Andalusia (Fig. 2).

**Fig. 2.**
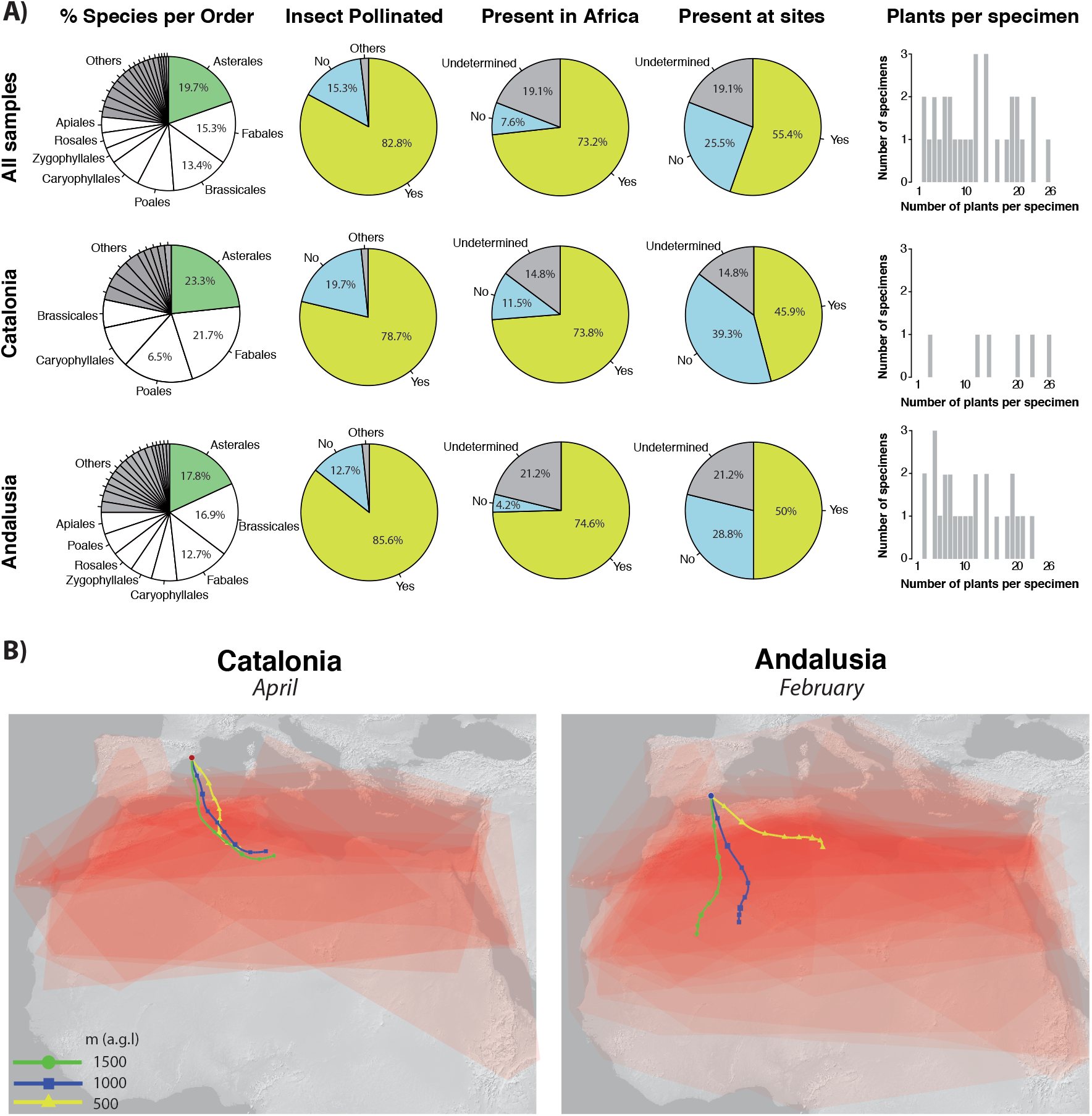
Detected plant species that are informative of migration: a) Pies showing classified percentages of plants per order, insect pollinated and presence in Africa and at sampling sites. Barplots show number of informative plants per specimen. Pies and barplots are shown for all samples together and for two migratory waves (Catalonia and Andalusia) independently. b) Additive extent of occurrences for informative plants detected in Catalonia and Andalusia migrants. Higher red intensity indicates higher probability of plant overlap). 48h backward wind trajectories at three altitudes (500m, 1000m and 1500 m agl) are shown (colour lines) for the specific dates of observed peaks of migration (April 27^th^ – Catalonia, February 21^st^ – Andalusia).

Sequences from all green plants (Viridiplantae) were present in the reference database and some sequences from our data matched green algae (Chlorophyta). Sequences from two samples were assigned to the algae genus of *Trebouxia.* Another species of green algae, *Pseudostichococcus monallantoides* was also detected in one sample.

Of the plant species detected from sequencing data, 40 were alien to the sampling sites (24 for Catalonia and 34 for Andalusia) and are thus potentially informative in estimating migratory paths (listed and illustrated in Tab. S3 and Fig. S4). From these, 33 species are present in Northern Africa and absent in Europe, and these were detected in 20 of our samples (Tab. S3). Six were present in more than two butterfly samples: *Calendula stellata* (9 butterfly samples) – occurring in Morocco, Algeria, Tunisia and Sicily, introduced in the Canary Islands; *Launaea mucronata* (6 samples) – a species of Saharo-Arabian distribution, also present in the Canary Islands; *Lotus weilleri* (4 samples) – a species endemic to northern Atlantic Morocco; *Oxalis compressa* (4 samples) – native to Southern Africa but introduced in the Mediterranean Region of Morocco and Algeria; *Raffenaldia primuloides* (4 samples) – present in Morocco and Algeria; *Farsetia aegyptia* (3 samples) – a desert species distributed throughout Northern Africa (Morocco, Algeria and Tunisia but absent from Libya and Egypt) and Asia (Tab. S3, Fig. S4).

## Discussion

### Pollen metabarcoding as a tool

Morphological identifications of pollen grains carried by insects have already been used to infer long-distance migratory patterns in insects (Mikkola, 1971; Hendrix et al., 1987; Hendrix & Showers, 1992; Gregg, 1993; Lingren et al., 1993, 1994; Westbrook et al., 1997). The use of this method, however, is limited because morphological identification by light microscopy is a time-consuming task, it requires specialized taxonomic expertise, and can hardly provide species-level determinations (Galimberti et al., 2014; Hawkins et al. 2015; Keller et al. 2015; Kraaijeveld et al. 2015; Richardson et al., 2015b; Sickel et al., 2015). DNA metabarcoding is a fast, high throughput method that greatly simplifies the identification process (Taberlet et al., 2012) and captures high diversity of pollen that is transported by insects. Our results show that identifying pollen grains carried by migrating insects through DNA metabarcoding is feasible to the species level and that high pollen diversity per specimen can be detected. Thanks to species-level identifications for multiple taxa, migratory paths of the insects can be traced and narrowed by additive geographic distributions of the plants. The method presented here has a wide application to all major insect orders that visit flowers (Coleoptera, Diptera, Hymenoptera and Lepidoptera; Kevan & Baker, 1983), as well as some vertebrates such as birds (Cronk & Ojeda, 2008) or bats (Fleming, Geiselman & Kress, 2009).

We show that the *V. cardui* individuals analysed here were migrating and originated in the African continent. Among the samples from which pollen was detected, all but three were carrying pollen attributed to plant species alien to their collecting site, most of it corresponding to African endemic plants. *Vanessa cardui* is typically colonizing the western Mediterranean Europe in spring, with most arrivals usually observed in April and May (Stefanescu et al., 2013). The origin of spring arrivals into the Iberian Peninsula has typically been associated to large breeding sources found in Morocco in March-April (Stefanescu, Alarcón, Izquierdo, Páramo & Ávila, 2007; Stefanescu, Páramo, Åkesson, Alarcón & Ávila, 2011). Our results partially agree with this timing and path. On the one hand, we show that arrivals to Europe can occur as early as February in Andalusia. On the other hand, the pollen associated to butterflies exhibits a much larger geographical range where butterflies might originate and transit (Fig 2). Unlike isotopic analyses, our approach does not test for natal origins of migrations, but for the most likely paths used during their migrations. During migration, butterflies generally stop in the evening, feed and rest at night. In the morning, they feed until it is warm enough and winds are suitable for continuing the migration (e.g., Shields, 1974). So, migratory paths would be defined by the position of stepping-stone locations where they fed. In fact, it is expected that pollen from most recently visited flowers is better represented on butterfly bodies than that from plants visited right after emergence. Thus, we may assume a “dilution effect” of the signal to a certain degree.

### Sequencing protocol

The flexible two-step laboratory protocol here presented can be easily adapted to other types of markers, for instance standard plant chloroplast barcodes such as rbcL and matK (CBOL Plant Working Group et al., 2009), just by replacing the primers in the first PCR reaction. Importantly, the proposed method allows for high sample multiplexing: up to 384 samples can be analysed simultaneously when combined with Illumina TruSeq CD index sequences, and even more when longer index sequences are used (e.g., Fadrosh et al., 2014). Previous pollen metabarcoding protocols used pollen pulverization with bullet blender and DNA isolation with commercial kits (Simel, Saidak & Tuskan, 1997; Kraaijeveld et al., 2015). Assessing the best pollen isolation method was beyond the scope of this study, but we found that skipping the pollen isolation step and using homogenized pollen directly in the PCR reaction with “direct” polymerase mix (Wong et al., 2014) is an efficient method of amplifying DNA markers from pollen loads. This way, DNA isolation – the most time-consuming step of pollen metabarcoding projects (Bell et al., 2017) – is avoided altogether. Nevertheless, further assessment of the effectiveness of such procedure in amplifying markers from all the plant species present in the pollen loads is necessary. Moreover, careful contamination and cross-contamination control by using blank samples and following best practices to avoid contamination are necessary in metabarcoding studies (Goldberg et al., 2016; Deiner et al., 2017). In our case, we used blank samples both at the pollen isolation and the library preparation steps. Both isolation and pre-PCR steps were also conducted under laminar flow cabinet. These remedial steps ensured no external contamination in our samples, as shown by blank samples and the virtual absence of plants native to Central Europe in our dataset.

Another source of bias are the errors occurring at the PCR and sequencing step (Coissac, Riaz, & Puillandre 2012). Many metabarcoding pipelines perform clustering of similar sequences in order to reduce the number of low-copy reads that are usually erroneous artefacts and cluster them together with the centroid sequence (e.g., Edgar, 2010; Rognes et al., 2016) or ‘denoising’ the reads in order to remove the putatively erroneous sequences (e.g., Edgar, 2016b). Studies with mock pollen samples are still needed to assess the relative performance of these methods for pollen metabarcoding.

Many factors can bias the PCR amplification of DNA templates and skew the quantitative representation of the sequenced species in the obtained reads. These factors involve PCR biases, either caused by sequence polymorphisms in priming sites (Sipos et al., 2007; Elbrecht & Leese, 2015; Piñol, Mir, Gomez-Polo & Agust, 2015), formation of chimeric reads (Bjørnsgaard Aas, Davey & Kauserud, 2017), sequence length or GC content (Krehenwinkel et al., 2017). Although some studies show some relationship between pollen abundance and the number of reads (Kraaijeveld et al., 2015; Pornon et al., 2016), other did not find such relationship (Hawkins et al., 2015; Richardson et al., 2015b). The number of reads obtained per species should be therefore treated with caution and only as a semi-quantitative method of estimating pollen abundance. Thus, following Yu et al. (2012), we used only presence-absence information when interpreting our results.

### Limitations to track migrations: taxonomic assignments and species distributions

Several key factors determine the accuracy and resolution of our method for studying insect migrations. First, reference sequences from correctly determined plant species are necessary to properly classify the obtained reads. Uneven geographical coverage of sequences present in reference databases, with a bias towards better studied areas such as Europe or Northern America (Ankerbrand, Keller, Wolf, Schultz & Förster, 2015; Bell et al., 2016), is of a special concern. Long-distance migration studies would benefit largely from global plant species coverage, which still remains a remote prospect. In our case, although we were able to classify many reads with high probability, a high proportion (52.6%) of unclassified reads remained. This is most likely caused by a number of African species/populations missing in our reference database and therefore classified with low probability, as much larger proportion of reads was classified into higher taxonomic ranks (Fig. S2). Representation of species in databases and taxonomic errors are especially problematic with best-hit approaches (e.g. BLAST; Boratyn et al., 2013), nevertheless such methods are still used in metabarcoding studies (Hawkins et al., 2015; Kraaijeveld et al., 2015; Richardson et al., 2015b). Sequence classifiers, such as SINTAX used here, are more robust in such cases. Nevertheless, all current methods display high over-classification rates in cases when taxa are missing from reference databases (Edgar, 2016a). So far, above classification methods have been mostly benchmarked for bacterial sequences (e.g., Vinje, Liland, Almøy & Snipen, 2015) and more studies are needed to assess their comparative performance for ITS2 metabarcoding of plant sequences.

In this study, we did not prioritize a full assessment of plant pollen present on the migrating butterflies and, in order to reduce false positives, we used a conservative 95% sequence assignment threshold for the classified reads to be retained in a final dataset. Despite such conservative approach, we could still detect a small number of species that are probably assignment errors, i.e. with geographical distributions outside the possible migratory routes of *Vanessa cardui.* This is probably because the representation of plant species belonging to taxonomically complex and diverse genera is far from complete in reference databases used for taxonomic assignments and could prevent the positive identification of some pollen grains. Some examples of such taxa in our dataset include: *Astragalus,* with about 2500–3000 species, is the largest genus of flowering plants (Podlech & Zarre, 2013; Bagheri, Maassoumi, Rahiminejad, Brassac & Blattner, 2017); *Artemisia* that covers approximately 600 species (Richardson, Page, Bajgain, Sanderson & Udall, 2012) and genus *Thymus* that includes around 400 species (Karaca, Ince, Aydin, Elmasulu & Turgut, 2015). Most of the species of these genera are native to the Mediterranean region, Northern Africa, and Western Asia. Moreover, molecular approaches have limitations to identify and define species in some of these complex genera due to various biological phenomena, such as interspecific hybridization and polyploidy, which are often correlated (Soltis & Soltis, 2009), can contribute significantly to the taxonomic complexity of *Thymus* (Morales, 2010), *Artemisia* (Richardson et al., 2012) and *Astragalus* (Doyle, 2012; Bagheri et al., 2017). In *Thymus*, genetic polymorphism at the intraspecific level can hinder the positive identification of some species (Karaca et al., 2015). Conversely, DNA sequence diversity is generally very low in some *Artemisia* species (Koloren, Koloren & Eker, 2016) and also among species included within several sections of *Astragalus* (Bagheri et al., 2017), which has been attributed to a rapid radiation (Sanderson & Wojciechowski, 1996).

In order to infer the migration routes from metabarcoding sequences, detailed plant distribution data are required. Evident gaps in the distribution of the plant species detected in North Africa exist. In particular, presence data available for Algeria and Libya are extremely poor when compared to Morocco or Tunisia. Such biases preclude more detailed analyses based on actual presence records or geographical grids (see Fig. S3 for results based on a presence grid). To avoid the influence of important gaps in presence data, distribution ranges delimited by peripheral presence records may be used (Fig. 2). The abovementioned limitations point to the importance of basic taxonomic, barcoding and floristic research, which is the cornerstone for myriad of studies.

More research is needed on pollen retention on insects. For instance, Del Scorro and Gregg (2001) found that the sunflower pollen is a transient marker that is only informative of plant visits that occurred during the previous two days. On the other hand, some studies, including the one presented here, show support for long-distance pollen transport (Hendrix & Showers, 1992; Ahmed et al., 2009). In this line, it is worth noting that no informative data was retrieved from a percentage of butterfly specimens (36%). In any case, pollen grains are probably lost along time, and a dilution effect of the pollen load signal is to be expected, with a higher representation of recently visited flowers.

Despite the abovementioned limitations, we have proven that pollen metabarcoding is an effective and informative method for tracking insect movements, despite the current limited resolution due to completeness of reference genetic libraries and plant species presence record databases. The accumulation of this kind of information grows rapidly thanks to the new sequencing techniques and citizen science initiatives, for example. As these two factors will most likely improve in the near future, the resolution and usefulness of pollen metabarcoding as a tool for tracking insect migrations can only increase.

### Pollen detected and migrations of *Vanessa cardui*

Using our metabarcoding approach, we were able to amplify a wide range of plant DNA sequences from migrating *Vanessa cardui.* We generally collected the butterflies immediately or soon after they landed on the beaches of the Mediterranean (note that in several instances we cannot discard that they fed on local flowers). Our study was therefore designed to test the feasibility of pollen detection after a long-distance migration. In this particular case, we expected to detect pollen of plant species present in Africa, including the Maghreb, the Sahara and the sub-Sahara. According to our hypothesis, large proportion of reads was classified as African or African-Arabian endemic plants (21.0%), or more generally, plants that were not native to the butterfly sampling sites (25.5%).

Pollen composition may explain individual migratory histories, but it can also report collective migratory histories given a time and site. Butterflies collected from the same spatiotemporal migratory waves (Andalusia in February and Catalonia in April) show parallelisms, but also differences, in their carrying pollen composition. Such differences among specimens could be explained by variability of the visited flowers and in the retention of pollen grains. In some cases, though, the plants found in different specimens are geographically exclusive, which suggest that, if taxonomic attributions are correct, the butterflies may have had different breeding origins and confluenced *a posteriori* during their migratory paths or at destination.

The two waves of migrants studied (Andalusia and Catalonia) could either correspond to populations originated during the winter in the Maghreb, or to populations originated in tropical Africa that may replenish the temperate zone in early spring (Talavera & Vila, 2016). The latter hypothesis cannot be excluded based on our data because, in addition to a marked influence from Maghreb flora, several floristic elements from the Sahara and Sub-Saharan Africa were detected (Fig. 2). Generally, the butterflies collected in February (Andalusia) showed a higher number of plants of predominantly Saharan distribution *(Farsetia stylosa, Launaea capitata, Launaea mucronata, Moltkioptis ciliata, Reseda villosa, Gymnocarpus decandrus, Euphorbia guyoniana),* or with an important sub-Saharan representation *(Pergularia tormentosa, Musa acuminata).* Pollen of *Musa* (banana), only detected in specimen 16C413, is extensively cultivated in tropical Africa and is abundant in the Canary Islands, but rarer in the Maghreb and Europe (where is generally cultivated in greenhouses).

In addition, this plant is apomictic and although some varieties preserve male flowers, these are very rare. As the probability to find male banana flowers is higher where the plant is common, a sub-Saharan origin for this sample is likely.

Butterflies from April (Catalonia) had few strictly Saharan floristic elements, such as *Launaea capitata* and *L. mucronata,* and mostly had representation of flora from the Maghreb (Fig. 2). Pollen endemic to the Canary Islands was detected in some instances, both from February and April. Thus, an origin in these islands cannot be discarded for some specimens.

Backtrack wind models agree with migratory paths coming from the African continent (Fig. 2), considering that *V. cardui* migrations can greatly be aided by winds (Stefanescu, Alarcón & Ávila, 2007). Winds consistently came from the south-east through central Algeria in both waves, at least during the previous 48h. Precise migratory sources or paths cannot be inferred based on pollen data or winds alone, but both can be combined to narrow predictions. Actually, the Algerian Maghreb (for the Catalonia April migratory wave) and the Algerian Sahara (for the Andalusia February wave) are the areas where higher accumulative probabilities of identified plants overlap. The combined evidence points to a highly probable origin or pass of these two migratory waves across Algerian grounds, although some individuals may have followed different paths.

Insect-pollinated plants prevailed in our results (82.8%), as expected in pollen obtained from butterfly bodies. Worth noting, plants detected belonged primarily to the Asterales, an order that includes many of *V. cardui* typical host plants and nectar sources (Nylin, Slove & Janz, 2014). In fact, the spectrum of plants visited by *V. cardui* is very wide, including most plant orders and life forms from small plants to trees. For example, numerous *V. cardui* specimens migrating northwards in February 2017 in south Morocco were observed to stop briefly to feed on the flowers of *Prunus dulcis* orchards (R. Vila, pers. obs.), a tree for which we detected pollen in seven butterfly specimens.

Most non-insect pollinated plants detected were wind-pollinated plants that belonged to the Poales. This points to the accidental transport of such pollen due to physical contact, as *V. cardui* tends to rest on the ground and grass, and only rarely on trees. Sequences from two samples were assigned to the algae genus *Trebouxia,* a common and widespread photobiont in lichens (Dal Grande et al., 2014). Lichens often reproduce asexually by soredia – small powdery propagules containing both fungus and algae. Such small structures could possibly stick to the body of butterflies accidentally and be transported on larger distances.

### Implications for plant pollination

Here we show that *Vanessa cardui* could potentially mediate transcontinental pollination, and thus gene flow, for plants species that occur in both Europe and Africa. Given the millions of individuals of *V. cardui* in particular, and of insects in general, that migrate every year across continents (Hu et al. 2016), the effects of this phenomenon on ecosystems and crops may be not negligible.

Regular long-distance gene flow is a phenomenon that has rarely been acknowledged, but that could potentially explain particular phylogeographic patterns in plants. For example, the Strait of Gibraltar has been shown as not effective at interrupting gene flow in *Androcymbium gramineum* (Caujapé-Castells & Jansen, 2003), several *Cistus* spp. (Fernández-Mazuecos & Vargas, 2010), *Hypochaeris salzmanniana* (Ortiz, Tremetsberger, Talavera, Stuessy & Garcia-Castano, 2007), and *Rosmarinus officinalis* (Mateu-Andrés et al., 2013). It is also worth noting that we detected sequences belonging to crops in our dataset – *Allium sativum, Cucumis sativus,* and *Prunus dulcis* – which points to the possibility of intercontinental pollination for these economically important species. We suggest that knowledge on the insect species that perform long-distance migration and the routes and temporal patterns they follow may be of high importance for better understanding intercontinental plant pollination, with implications extending from plant phylogeography to ecosystem services.

**Table 1.**
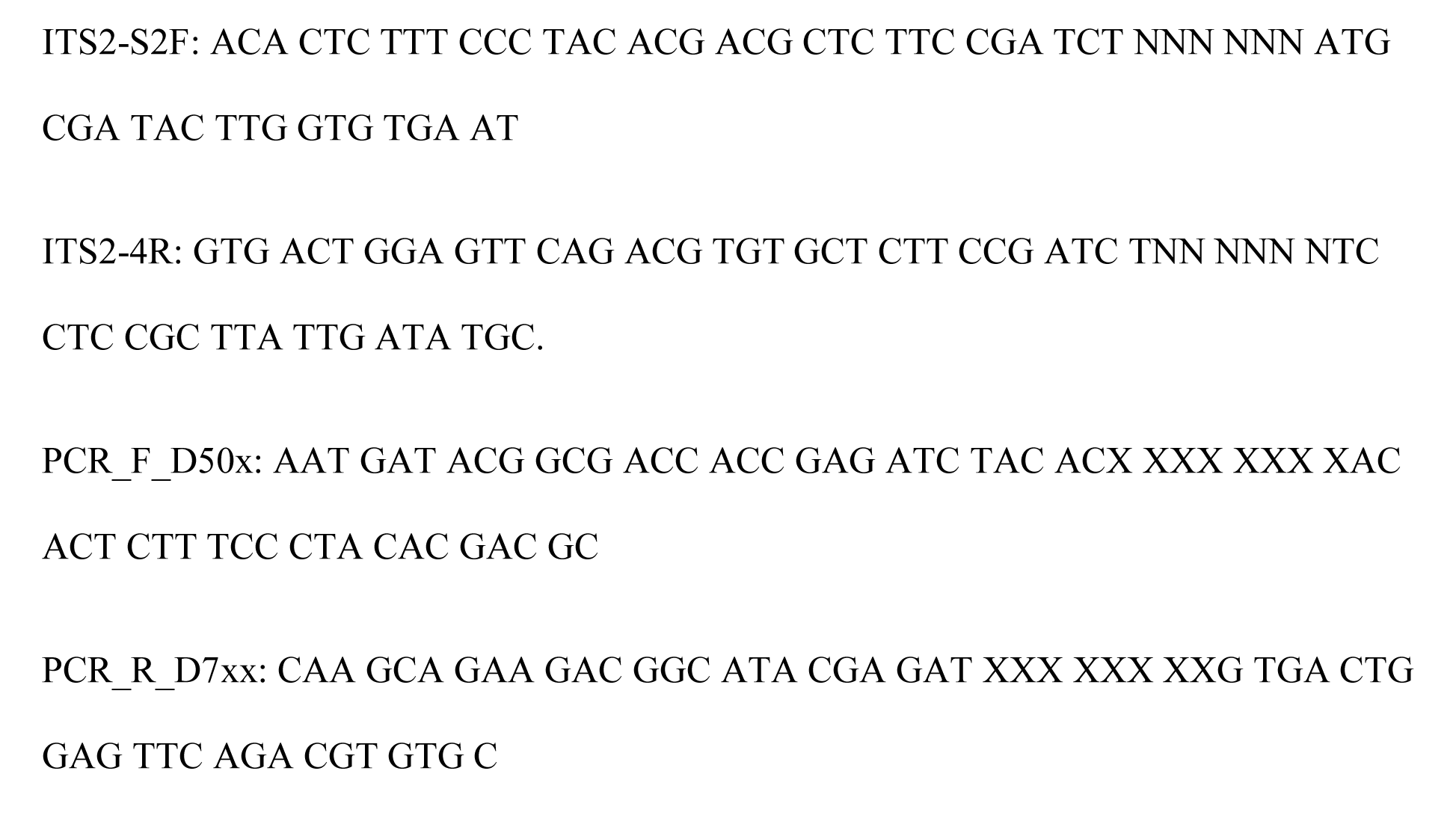
Sequences of the primers used: ITS2-S2F and ITS2-4R primers were used in the first PCR reaction; PCR_F_D50x and PCR_R_D7xx are the indexing primers used in the second PCR reaction; XXX XXX XX are 8 nt-long index sequences.

## Acknowledgements

We thank Martin Gascoigne-Pees for important sample contribution. TS and MR are supported by the statutory funds of W. Szafer Institute of Botany, Polish Academy of Sciences. GT is supported by the Marie Curie Actions FP7-People-2013 IOF (project 622716) and by the MINECO programme Juan de la Cierva Incorporación (IJCI-2016-29083).

Funding was provided by the British Ecological Society (grant LRB16/1015) to GT, by project CGL2016-76322 (AEI/FEDER, UE) and by the Committee for Research and Exploration of the National Geographic Society (grant WW1-300R-18) to GT and RV.

## Data accessibility

Scripts used to process the data: https://github.com/TomaszSuchan/pollen-metabarcoding

ITS2 database for SINTAX: https://github.com/molbiodiv/meta-barcoding-dual-indexing/blob/master/precomputed/viridiplantae_all_2014.sintax.fa

Raw sequences: European Nucleotide Archive, http://www.ebi.ac.uk/ena/data/view/PRJEB26439

## Author contributions

TS, GT and RV conceived the study and wrote the manuscript. GT and RV collected samples. TS carried out laboratory work. TS and GT analyzed data. LS gathered plant distribution data. All authors edited and approved the final version of the manuscript.

